# Fluid Shear Stress-Induced Changes in Megalin Trafficking Enhance Endocytic Capacity in Proximal Tubule Cells

**DOI:** 10.1101/2024.02.22.581213

**Authors:** Emily M. Lackner, Isabella A. Cowan, Kimberly R. Long, Ora A. Weisz, Katherine E. Shipman

## Abstract

Proximal tubule (PT) cells maintain a high-capacity apical endocytic pathway to recover essentially all proteins that escape the glomerular filtration barrier. The multiligand receptors megalin and cubilin play pivotal roles in the endocytic uptake of normally filtered proteins in PT cells but also contribute to the uptake of nephrotoxic drugs, including aminoglycosides. We previously demonstrated that opossum kidney (OK) cells cultured under continuous fluid shear stress (FSS) are superior to cells cultured under static conditions in recapitulating essential functional properties of PT cells *in vivo*. To identify drivers of the high-capacity, efficient endocytic pathway in the PT, we compared FSS-cultured OK cells with less endocytically active static-cultured OK cells. Megalin and cubilin expression are increased, and endocytic uptake of albumin in FSS-cultured cells is >5-fold higher compared with cells cultured under static conditions. To understand how differences in receptor expression, distribution, and trafficking rates contribute to increased uptake, we used biochemical, morphological, and mathematical modeling approaches to compare megalin traffic in FSS-versus static-cultured OK cells. Our model predicts that culturing cells under FSS increases the rates of all steps in megalin trafficking. Importantly, the model explains why, despite seemingly counterintuitive observations (a reduced fraction of megalin at the cell surface, higher colocalization with lysosomes, and a shorter half-life of surface-tagged megalin in FSS-cultured cells), uptake of albumin is dramatically increased compared with static-grown cells. We also show that FSS-cultured OK cells more accurately exhibit the mechanisms that mediate uptake of nephrotoxic drugs *in vivo* compared with static-grown cells. This culture model thus provides a useful platform to understand drug uptake mechanisms, with implications for developing interventions in nephrotoxic injury prevention.

## 1 Introduction

The proximal tubule (PT) is comprised of polarized epithelial cells that are responsible for recovering essentially all proteins that escape the glomerular filtration barrier by a specialized, high-capacity endocytic pathway to prevent protein loss in the urine (Weisz, 2021; Eshbach and Weisz, 2017). Receptors are internalized via clathrin-coated pits from apical membrane invaginations (Hatae et al., 1986; Eshbach and Weisz, 2017). Once clathrin is dissociated, these vesicles fuse with apical early endosomes (AEEs), and then quickly acidify and mature into apical vacuoles (AVs) (Shi et al., 1991; Hatae et al., 1986). Recycling of receptors occurs from AEEs and AVs via dense apical tubules (DATs) that fuse with the apical membrane (Nielsen, 1993). Ligands are targeted to the lysosomes for degradation (Weisz, 2021; Eshbach and Weisz, 2017; Christensen et al., 2012).

Protein recovery by PT cells is mediated by the abundantly expressed large multiligand receptors megalin and cubilin (Christensen et al., 2012; Moestrup and Verroust, 2001). Megalin (>600 kDa) contains four ligand binding sites and a transmembrane domain with a cytoplasmic tail that contains binding motifs for Dab2, a clathrin-adaptor protein (Beenken et al., 2023; Eshbach and Weisz, 2017). Cubilin (>440 kDa), which does not contain a transmembrane domain, associates as a trimer with amnionless to form the cubilin-amnionless (CUBAM) complex (Weisz, 2021; Eshbach and Weisz, 2017; Larsen et al., 2018). Amnionless has a transmembrane domain and cytoplasmic tail, which contains binding motifs for Dab2 (Ahuja et al., 2008; Larsen et al., 2018). While megalin and CUBAM physically interact and are believed to traffic together in the PT, they can function independently as well (Rbaibi et al., 2023; Shipman et al., 2022; Long et al., 2022; Park et al., 2020).

Megalin and cubilin bind a number of ligands with varying affinity (Moestrup and Verroust, 2001; Nielsen et al., 2016; Zhai et al., 2000). For example, cubilin serves as a high affinity receptor that recovers normally-filtered levels of albumin, while megalin binds to albumin with low affinity and provides additional capacity when tubular albumin levels are increased (Edwards et al., 2022; Rbaibi et al., 2023; Ren et al., 2020; Weisz, 2021). In addition, many clinically important drugs, including the aminoglycoside antibiotic gentamicin, bind directly to megalin. Knockout or inhibition of megalin reduces their toxicity in rodents and in L2 carcinoma cells (Nagai and Takano, 2014; Wagner et al., 2023; Nagai et al., 2001; Moestrup et al., 1995; Quiros et al., 2011). Strikingly, we recently showed that loss of megalin expression or function not only decreases receptor-mediated endocytic uptake, but also greatly impairs the volume of membrane and fluid (flux) moving through endocytic compartments (Rbaibi et al., 2023; Long et al., 2022). This occurred in the absence of any changes in the expression of endocytic compartment markers or their distribution (Rbaibi et al., 2023). Thus, megalin expression can function as a positive driver of all endocytic traffic, beyond its own ligands.

Previously, our lab showed that opossum kidney (OK) cells cultured under continuous fluid shear stress (FSS) develop a high-capacity endocytic pathway that is similar in organization and morphology to that of the PT *in vivo* (Long et al., 2017; Ren et al., 2019; Park et al., 2020; Long et al., 2022). In addition, OK cells cultured under these conditions exhibit higher expression of megalin and cubilin, greater endocytic capacity, more apical microvilli and basolateral invaginations enfolding mitochondria, more lysosomes, and greater utilization of oxidative phosphorylation to drive metabolic needs compared to cells cultured under static conditions (Long et al., 2017; Ren et al., 2019). We recently developed kinetic models that describe megalin trafficking in FSS-cultured OK cells and in a genetic model of Dent disease that allowed us to pinpoint specific trafficking steps impacted by loss of ClC-5 (Shipman et al., 2022, 2023). These models enable a better understanding of the mechanisms that drive endocytosis, as well as drug-induced nephrotoxicity. To identify key drivers of the efficient, high-capacity endocytic pathway of the PT, we utilized our cell culture model system and adapted our megalin trafficking mathematical model to understand the basis for increased endocytic capacity in cells cultured under FSS.

## 2 Materials and Methods

### 2.1 Cell Culture

OK cells were cultured in DMEM-F12 (Sigma; D6421) supplemented with 2.5x GlutaMax (Gibco; 35050-061) and 5% fetal bovine serum (FBS) at 37°C and 5% CO_2_. For experiments, cells were seeded on 12 mm Transwell permeable supports (Costar, 3401) in 12-well dishes at 4x10^5^ cells per 0.5 mL medium on the apical side of the filter. The basolateral side of the filter received 1.5 mL of medium. After overnight incubation, the filters were transferred to an orbital platform shaker in the incubator and rotated at 146 rpm for 72 h to enhance differentiation (FSS) or maintained under static conditions, as described previously (Long et al., 2017). Media was changed daily.

### 2.2 Quantitation of uptake

#### 2.2.1 Albumin and dextran

Cells cultured on filters under static or FSS conditions as described above were washed in DMEM-F12, 2.5x GlutaMax, and 25mM HEPES (Gibco, 15630-080; DF+H Media) and incubated at 37°C under continued 1X or 0X conditions for 15-30 minutes as indicated with 40 µg/mL AlexaFluor-647 albumin (Invitrogen, A34785) or 0.5 mg/mL AlexaFluor-647 10-kDa dextran (Invitrogen, D22914) added apically in DF+H medium. Filters were washed three times with cold phosphate-buffered saline containing MgCl_2_ and CaCl_2_ (PBS; Sigma, D8662), filters were excised with a razor blade, and cells were solubilized in 300 µL detergent lysis buffer (50 mM Tris, pH 8.0, 62.5 mM EDTA, 1% IGEPAL, 4 mg/ml deoxycholate) for 30 minutes on a rotating platform rocker at 4°C, as described previously (Shipman et al., 2023). Uptake of albumin or dextran fluorescence was quantified by spectrofluorimetry. Uptake was normalized to account for the 1.3-fold increase in cell density in FSS-cultured cells compared with static-grown cells (Long et al., 2017).

#### 2.2.2 Gentamicin

Cells cultured on filters under static or FSS conditions as described above were washed in DMEM-F12, 2.5x GlutaMax, and 25mM HEPES (Gibco, 15630-080; DF+H Media) and incubated at 37°C under continued static or FSS conditions for 3 hours with 50-1000 nM Texas Red gentamicin conjugate (GTTR; AAT Bioquest, 24300) apically in DF+H medium. The filters were then lysed, and uptake was measured following the same protocol as albumin uptake.

#### 2.2.3 Uptake in the presence of RAP

Cells were pretreated for 30 min at 37°C with 0.5mM of apically added human RAP (Innovative Research, IHURAP) in DF+H medium. Alexa Fluor-647 albumin (40 µg/mL), AlexaFluor-647 dextran (0.5 mg/mL), or GTTR (500 nM) was added apically in the continued presence RAP and uptake was quantified as described above.

### 2.3 Surface binding of albumin

Cells cultured on filters were incubated for 30 min on ice with 100 µg/mL AlexaFluor-647 albumin added apically in DF+H media. Filters were washed 3x rapidly in ice-cold PBS, excised with a razor blade, and cells solubilized in 300 µL detergent lysis buffer for 30 min on a rotating platform rocker at 4°C. Surface-bound AlexaFluor-647 albumin was quantified by spectrofluorimetry. Surface binding was normalized to determine binding per cell (Long et al., 2017).

### 2.4 Western blotting of cell lysates

Filters were washed with cold PBS, excised, and solubilized in 300 µL detergent lysis buffer containing 5 µg/mL leupeptin, 7 µg/mL pepstatin A, 1M PMSF, and a Pierce Protease Inhibitor Mini Tablet (Thermo Scientific, A32955; 1 tablet/10 ml of buffer). for 20 min on a rotator at 4°C. The protein concentration of the lysate was measured using a DC Protein Assay Kit (Bio-Rad; 5000111). Equivalent amounts of total protein were separated by SDS-PAGE on 4% to 15% Criterion TGX gels (Bio-Rad; 5671083). Megalin and cubilin were detected with anti-megalin antibody generously provided by Dr. Daniel Biemesderfer and Dr. Peter Aronson (Yale University, MC-220, 1:20,000 (Zou et al., 2004)) and polyclonal anti-cubilin antibody (No. 27445, validated in (Ren et al., 2020; Long et al., 2022) 1:5000), respectively. Cells cultured under FSS yielded 2.1-fold more protein/filter compared with cells cultured under static conditions. Data were normalized to determine relative receptor expression per cell based on the higher protein levels in FSS-versus static-grown cells.

### 2.5 Surface biotinylation based assays

#### 2.5.1 Endocytosis

OK cells cultured on permeable supports under static or FSS conditions were washed with cold PBS and the apical surface biotinylated with 1 mg/mL EZ-Link Sulfo-NHS-SS-biotin (Thermo Scientific, 21331) in 0.5 mL TEA-buffered saline (TBS; 10 mM triethanolamine-HCl, pH 7.6, 137 mM NaCl, 1 mM CaCl_2_) for 2x 15 min on ice. The biotinylation reaction was quenched by washing with DMEM-F12 plus 5% FBS for 10 min on ice. Samples were rinsed once with ice cold DF+H media then quickly warmed to 37°C by the addition of prewarmed DF+H and placed in the incubator either on an orbital shaker (FSS) or left under static conditions for 0-5 min. Endocytosis was stopped by washing with prechilled PBS on ice. Biotin at the cell surface was stripped by washing cells with prechilled 100 mM MESNA in Stripping Buffer (50 mM Tris-HCl pH 8.6, 100 mM NaCl, 1 mM EDTA, 0.2% BSA) for 2x 20 min on ice. A duplicate 0 min time-point was left unstripped to quantify the fraction of total megalin present at the apical surface at steady state. Residual MESNA was quenched by washing cells with ice cold DF+H media for 10 min on ice. Filters were washed with ice-cold PBS, excised with a clean razor blade, and solubilized in 600 µL detergent lysis buffer for 20 min at 37°C. To determine total megalin levels, 5% of the lysate volume was reserved. Biotinylated proteins were precipitated from the remaining lysate by overnight incubation at 4°C with streptavidin agarose resin (Thermo Scientific, #20353) and recovered in 4x loading sample buffer (0.2M Tris-HCl pH 6.8, 8.6M glycerol, 8% SDS, 0.025% bromophenol blue) with 5% 2-mercaptoethanol by heating at 98°C for 5 min. Samples were analyzed by Western blot after SDS-PAGE on 4-15% Criterion TGX gels.

#### 2.5.2 Half-life of surface-biotinylated megalin

The apical surface of filter-grown OK cells was biotinylated as above. Cells were rinsed once with ice cold DF+H media then quickly warmed to 37°C by the addition of prewarmed DMEM-F12 plus 5% FBS and placed in the incubator either under static or FSS conditions. At each time point starting from 1 to 6 h, filters were rinsed in cold PBS, cells were lysed, biotinylated proteins were recovered, and samples were blotted to quantify remaining biotinylated megalin as described above.

### 2.6 Immunofluorescence staining

Fluorescence detection and quantitation of the colocalization between megalin and endocytic compartment markers was performed as described in Shipman et al. 2022 (Shipman et al., 2022). Briefly, filters were washed in warm PBS and fixed in warm 4% paraformaldehyde (PFA) and 100mM sodium cacodylate at ambient temperature. After 2 washes in PBS, filters were quenched (PBS, 20mM glycine, and 75mM ammonium chloride) for 5 min and permeabilized for 5 min in quench solution containing 0.1% Triton X-100. After washing with PBS, the filters were blocked with PBS, 1% BSA, and 0.1% saponin, and incubated for 1 h with primary antibody diluted in PBS, 0.5% BSA, and 0.025% saponin (wash buffer). The filters were washed 3 times, incubated for 30 min with secondary antibody diluted in wash buffer, and washed 3 times. After excising, filters were mounted onto glass slides with ProLong Glass Antifade Mountant (Molecular Probes, P36981). For costaining with primary antibodies from different host species, filters were incubated with both primary antibodies simultaneously followed by both secondary antibodies (Negoescu et al., 1994). When costaining with primary antibodies from the same host species labeling was performed sequentially, as previously described in Shipman et al. 2023. Antibodies, sources, and dilutions used for indirect immunofluorescence in OK cells are listed in Table S1.

Filters were imaged on a Leica Stellaris-8 inverted confocal microscope using a 63x oil immersion objective (NA 1.4). Filters stained with Cathepsin B were imaged on the Leica SP8 confocal microscope with a 63x oil objective (NA 1.4). All images were acquired with a voxel size of 45x45x130 nm (x, y, z). All images were deconvolved with Huygens Essential version 17.04 using the CMLE algorithm, with SNR 20 and 40 iterations (Scientific Volume Imaging, The Netherlands, http://svi.nl). Colocalization of two channels over the whole z-stack was determined by Manders’ coefficients using the JACoP plugin for ImageJ without thresholding (Bolte and Cordelières, 2006; Schneider et al., 2012).

### 2.7 Mathematical model of megalin trafficking

We adapted our previously described comprehensive model of megalin traffic in FSS-cultured OK cells (Shipman et al., 2022) to predict kinetic trafficking rates for cells cultured under static and continuous FSS conditions. The model describes the traffic of megalin from the apical surface through intracellular compartments of PT cells. Kinetic data from biotinylation-based assays and colocalization data collected from static- and FSS-cultured cells were used in the estimation of kinetic parameters as previously described (Shipman et al., 2022) with minor modifications described below.

#### 2.7.1 Endocytic Rate

The biotinylated megalin remaining at the surface during the brief endocytosis period can be represented as:

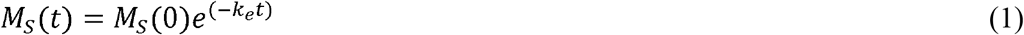

where *M*_*S*_(*t*) is the biotinylated megalin remaining at the surface at time *t* and *M*_*S*_(0) = 100. The data obtained from the endocytosis biotinylation assay represents the percent of surface megalin that is internalized over time and is equivalent to *M*_*int*_(*t*) = *M*_*S*_(0) − *M*_*S*_(*t*). To estimate the endocytic rate of megalin, the data from 0-5 min were log transformed and fit with a simple linear regression model such that:

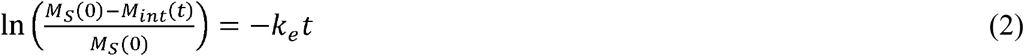

The slopes of the lines of best fit for static- and FSS-cultured cells were compared with the extra sum-of-squares F-test to see if they were significantly different. With a p-value = 0.0012, it was determined that the slopes were significantly different. Statistical analysis was performed in GraphPad Prism 9.3.1.

#### 2.7.2 Intracellular Trafficking Rates

The steady state distribution of megalin among endocytic compartments was calculated using the megalin colocalization values in Table S2 and the colocalization values for the overlap between different combinations of endocytic markers in Table S3 as described previously. However, we used Cathepsin B staining as the marker for lysosomes instead of LysoTracker Red DND-99. Standard error was propagated accordingly. This steady state distribution was used to estimate intracellular kinetic rates with the assumption that 51% of recycling megalin traffics through AVs (*α = 0*.*51*) for all cell lines (Shipman et al., 2022).

## 3 Results

### 3.1 Cells cultured under continuous shear stress have increased endocytic capacity and total protein

To establish the differences in endocytic capacity and receptor expression in cells grown under either static or continuous FSS conditions, we profiled the difference in endocytic uptake and receptor expression per cell. Albumin at normally filtered concentrations (40 µg/mL) binds primarily (approximately 90%) to cubilin (Ren et al., 2020), whereas megalin expression drives the overall rate of uptake (Long et al., 2023). FSS-cultured cells had 5.4-fold higher endocytic uptake of albumin per cell (Figure 1A) and about 2.8-fold higher uptake of dextran per cell (a marker of the fluid that accompanies albumin into endocytosed vesicles) compared with cells cultured under static-conditions (Figure 1B). By contrast, there is only a 3.8-fold increase in megalin expression and 2.3-fold increase in cubilin expression per cell in FSS-cultured cells (Figure 1C-D). The increase in receptor expression, therefore, does not fully account for the total increase in endocytic capacity in FSS-cultured cells. Thus, we looked to see if there was also a difference in receptor localization in FSS-versus static-cultured cells.

**Figure 1.**
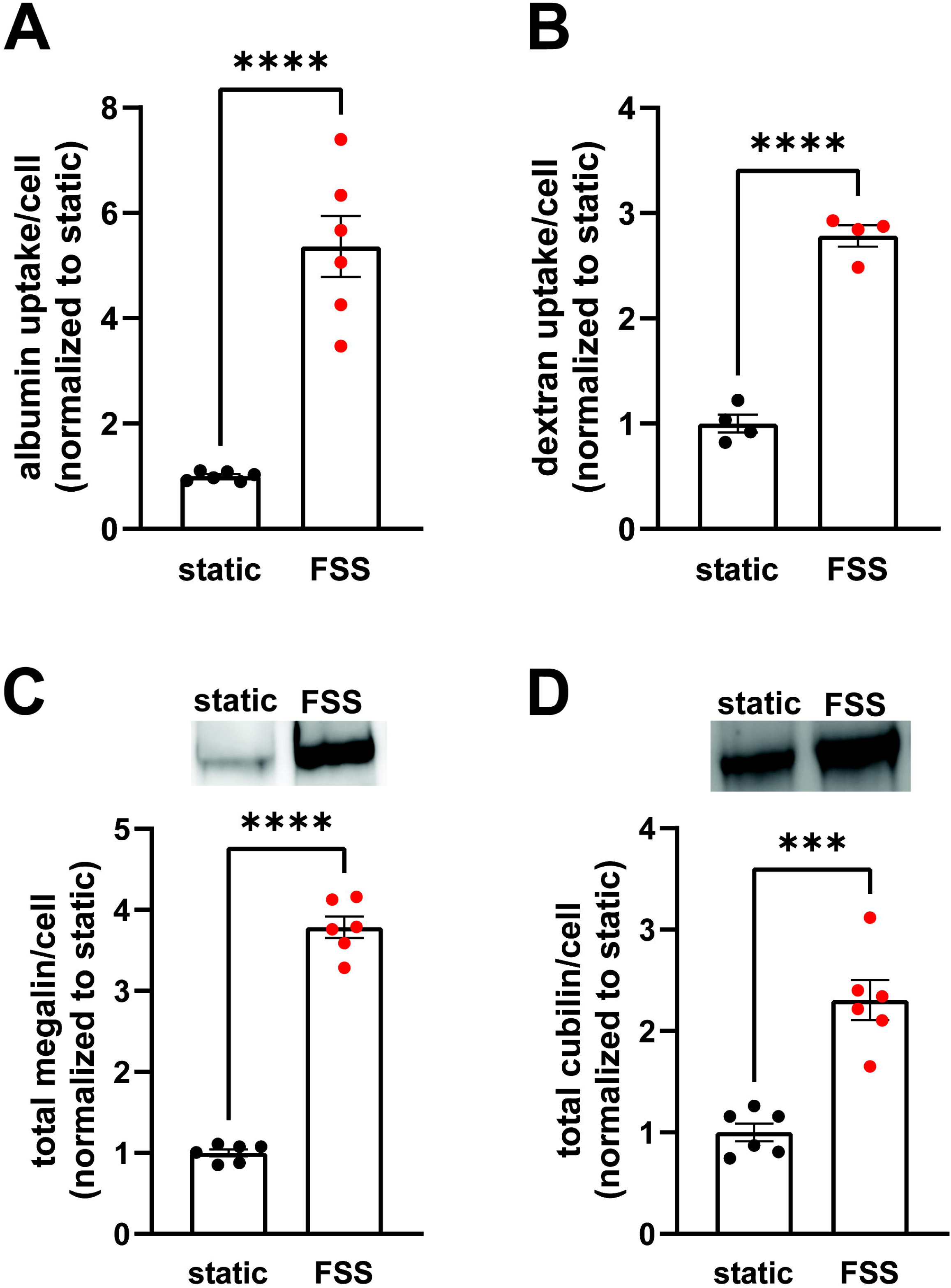
OK cells have increased expression of megalin and cubilin and greater endocytic capacity when cultured under continuous shear stress. (A) OK cells were incubated for 15 minutes with 40 µg/mL AlexaFluor-647 albumin and then solubilized. Fluorescent albumin uptake of cells cultured under static conditions or under continuous fluid shear stress (FSS) was quantified by spectrofluorimetry per cell. Data from 3 independent experiments done in duplicate were normalized to static OK albumin uptake. ****P=<0.0001 by unpaired t-test. (B) OK cells were incubated for 30 minutes with 0.5mg/mL AlexaFluor-647 dextran and then solubilized. Fluorescent dextran uptake was quantified by spectrofluorimetry per cell. Data from 1 experiment with 4 replicates were normalized to static OK dextran uptake. ****P=<0.0001 by unpaired t-test. (C-D) Equivalent amounts of protein from lysates of static-cultured and FSS-cultured cells were blotted for megalin (C) and cubilin (D). The quantified band intensity of 6 replicates was normalized to the total average of static-cultured cells. The data was transformed to fold-change in expression rate per cell by accounting for the 2.1-fold increase in total protein per filter and 1.3-fold increase in cell density of FSS-cultured cells. Representative blots are shown above each graph. ****P=<0.0001, ***P=0.0001 by unpaired t-test.

### 3.2 Biochemical analysis of receptor localization and traffic

Because the increase receptor expression does not fully explain the increase in endocytic capacity, we compared receptor localization by quantifying both ligand binding sites and the fraction of total receptors located at the apical surface. FSS-cultured cells had a numerically greater number of albumin surface binding sites per cell, although not statistically significant, compared to static-cultured cells (Figure 2A). Interestingly, surface biotinylation of megalin revealed a 43% decrease in the fraction of total megalin at the apical membrane of FSS-cultured cells (2.8%) when compared to static-cultured cells (4.9%; Figure 2B). The fraction of total cubilin at the surface decreased by ~60% in FSS-cultured cells (6.6%) when compared to static-cells (16.2%; Figure 2C). Interestingly, the fraction of both megalin and cubilin at the surface in FSS-cultured cells is more similar, (2.8% megalin, 6.5% cubilin) than in static-cultured cells (4.9% megalin, 16.2% cubilin). These changes in receptor surface localization suggest that there are differences in trafficking kinetics in addition to the increase in the total number of the receptors in FSS-cultured cells.

**Figure 2.**
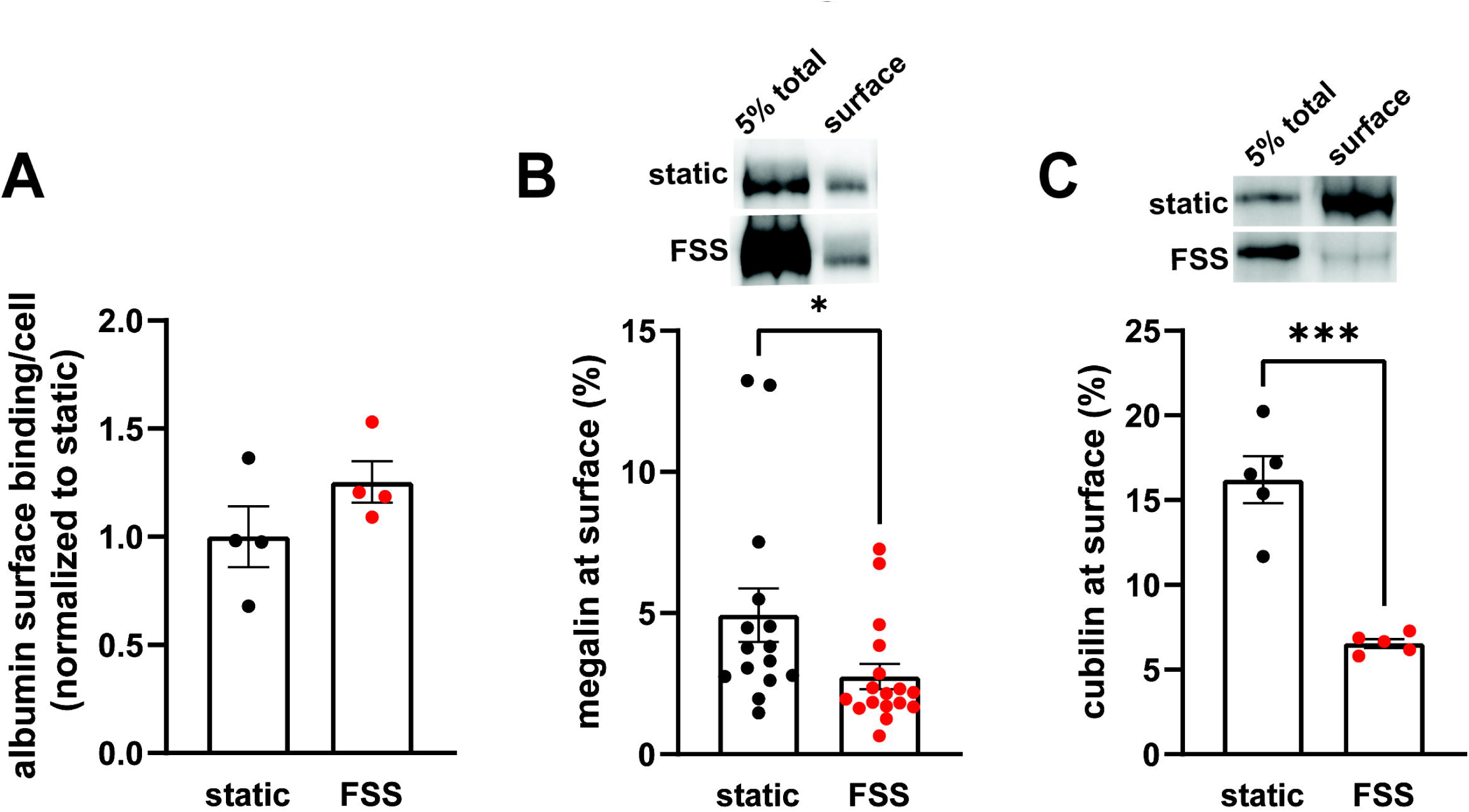
FSS-cultured cells have a greater number of binding sites but a reduced fraction of total receptors at the surface. (A) OK cells were incubated on ice for 30 minutes with 100 µg/mL AlexaFluor-647 albumin and then solubilized. Surface bound albumin was quantified by spectrofluorimetry per cell. Data from 1 experiment done in quadruplicate were normalized by experiment to the static OK cell surface binding. P=0.1868 by unpaired t-test. (B-C) The apical surface of OK cells was biotinylated on ice and cells were then lysed. Biotinylated megalin (B) and cubilin (C) were recovered using streptavidin beads and western blotted using anticubilin and antimegalin antibody, respectively. Surface cubilin and megalin were quantified as a percentage of total cubilin/megalin for each condition. Data from 6-11 independent experiments are plotted. ***P=0.0001, *P=0.0390 by unpaired t-test. Representative blots of the total (5%) and surface cubilin and megalin in static-cultured and FSS-cultured OK cells are shown above the graphs.

Because megalin drives endocytic flux, we compared megalin trafficking rates in cells cultured under static versus FSS conditions (Long et al., 2023). The endocytic rate of megalin in cells cultured under static and FSS conditions was quantified by measuring biotinylated surface megalin internalized over a five minute interval and normalized to surface megalin. FSS-cultured cells have a 3.1-fold increase in endocytic rate compared to static-cultured cells (Figure 3A; P=0.0012, extra sum of squares F-test). The increased endocytic rate in FSS-cultured cells is consistent with the lower steady state fraction of receptors at the apical surface under these conditions. Taking into account the levels of total and surface megalin, we calculate a 6.8-fold increase in the number of megalin molecules internalized per minute per cell in FSS-cultured cells compared with static-grown cells. To determine whether subsequent trafficking steps are also affected by culture under continuous shear stress, we quantified the half-life of surface-biotinylated megalin. The half-life was about 34% shorter in FSS-cultured cells than in static-cultured cells (2.6h vs. 3.9h, respectively), consistent with an increase in endocytic rate (Figure 3B).

**Figure 3.**
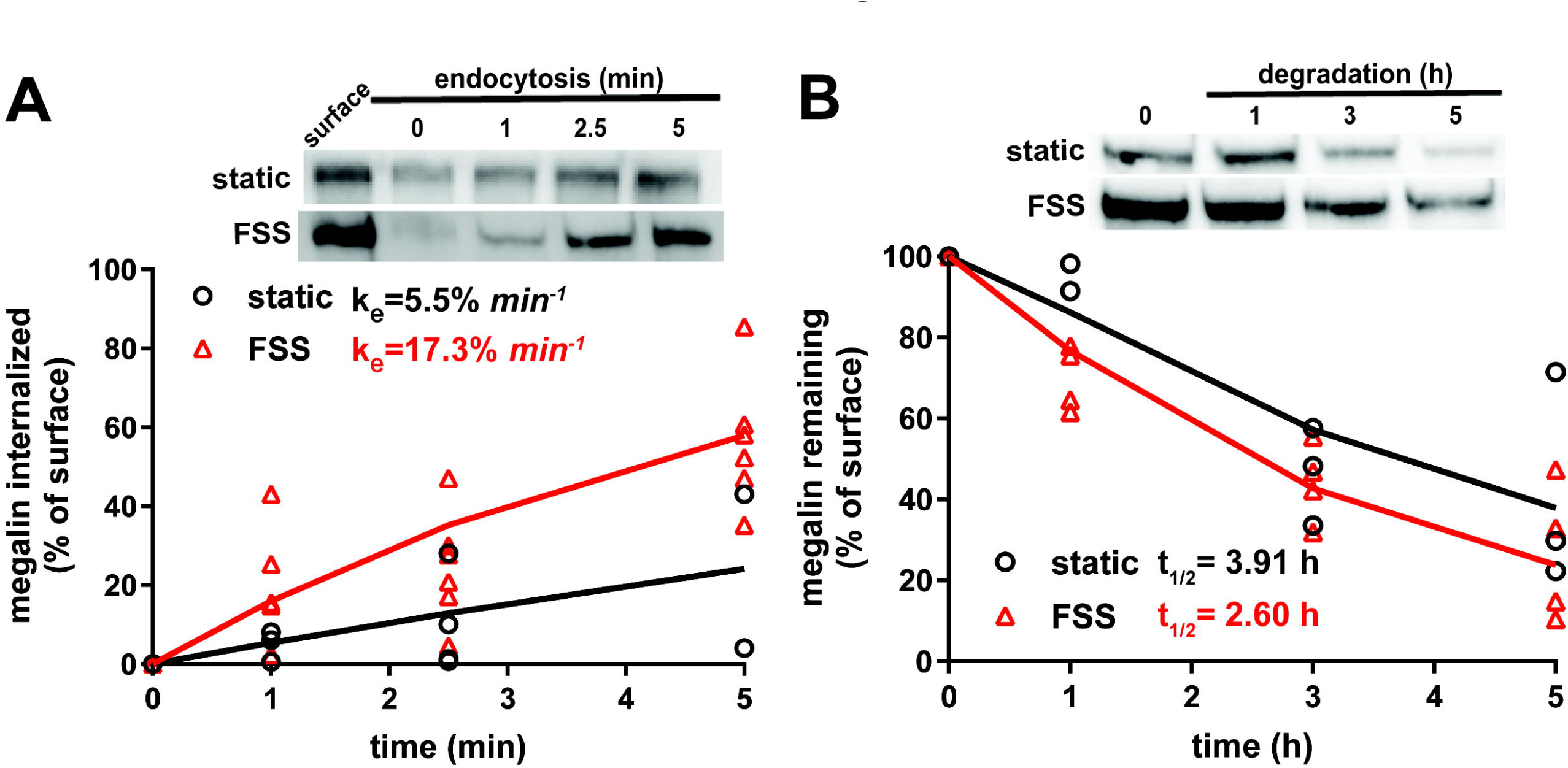
Biochemical measurements of megalin traffic show an increased rate of trafficking in FSS-cultured cells. (A) The apical surface of OK cells was biotinylated on ice and returned to culture for 0-5 min. Biotin remaining at the apical surface was stripped at each time point. Cells were lysed and biotinylated megalin was recovered with streptavidin beads and was blotted. Internalized megalin was quantified as a percentage of surface megalin at steady state for each condition. Data from four static-cultured (black) and eight FSS-cultured (red) independent experiments are plotted. The data were transformed and fit to a line to determine the fractional endocytic rate as previously described (Shipman et al., 2022). Lines represent predicted values. The endocytic rate as a percent of surface megalin for static and FSS-cultured cells is shown on the top left of the graph. P=0.0012 by extra sum of squares F-test. The estimated endocytic rates as percent of surface megalin internalized per minute are shown in the top left corner of the graph. Representative blots are shown above the graph. (B) The degradation of apically biotinylated megalin was determined by returning cells to culture for extended time periods. At each time point, the cells were lysed and recovered biotinylated megalin was blotted and quantified as a percentage of T=0h for each condition. Data from 3 static-cultured (black) and 4 FSS-cultured (red) independent experiments are plotted and were used to fit (lines) the degradation rate of megalin using estimates of fractional endocytic rate and the fraction of megalin at the apical surface (Figure 2C). The estimated half-life of surface megalin in static-cultured versus FSS-cultured cells is shown in the bottom left corner of the graph. Representative blots are shown above the graph.

### 3.3 Mathematical model reveals differences in megalin trafficking in cells cultured under static or shear stress conditions

We used quantitative imaging to determine the steady state colocalization of megalin in cells cultured under static or FSS conditions with respect to known markers of apical endocytic compartments. Cells were co-stained to detect megalin and markers for AEEs (EEA1), AVs (Rab7), DATs (Rab11a), and lysosomes (Lys; Cathepsin B), as shown in Figure 4A. Cells cultured under static conditions are markedly shorter than FSS-cultured cells (Figure S1), however, the markers we looked at were similarly distributed except for Rab11a, which was localized subapically in FSS-cultured cells but had a more dispersed distribution in static-cultured cells (Figure 4D). The colocalization of megalin with each marker was quantified by Manders’ coefficient. Colocalization of megalin with EEA1, Rab7, and Rab11a was quantitatively similar in static- and FSS-cultured cells (Figure 4B-E, Table S2). Colocalization of megalin with Cathepsin B was greater in FSS-cultured cells (Figure 4B, F, Table S2), which is consistent with the increased staining of Cathepsin B in the cells, as well as increased lysosome number in FSS-cultured cells as previously reported (Long et al., 2017).

**Figure 4.**
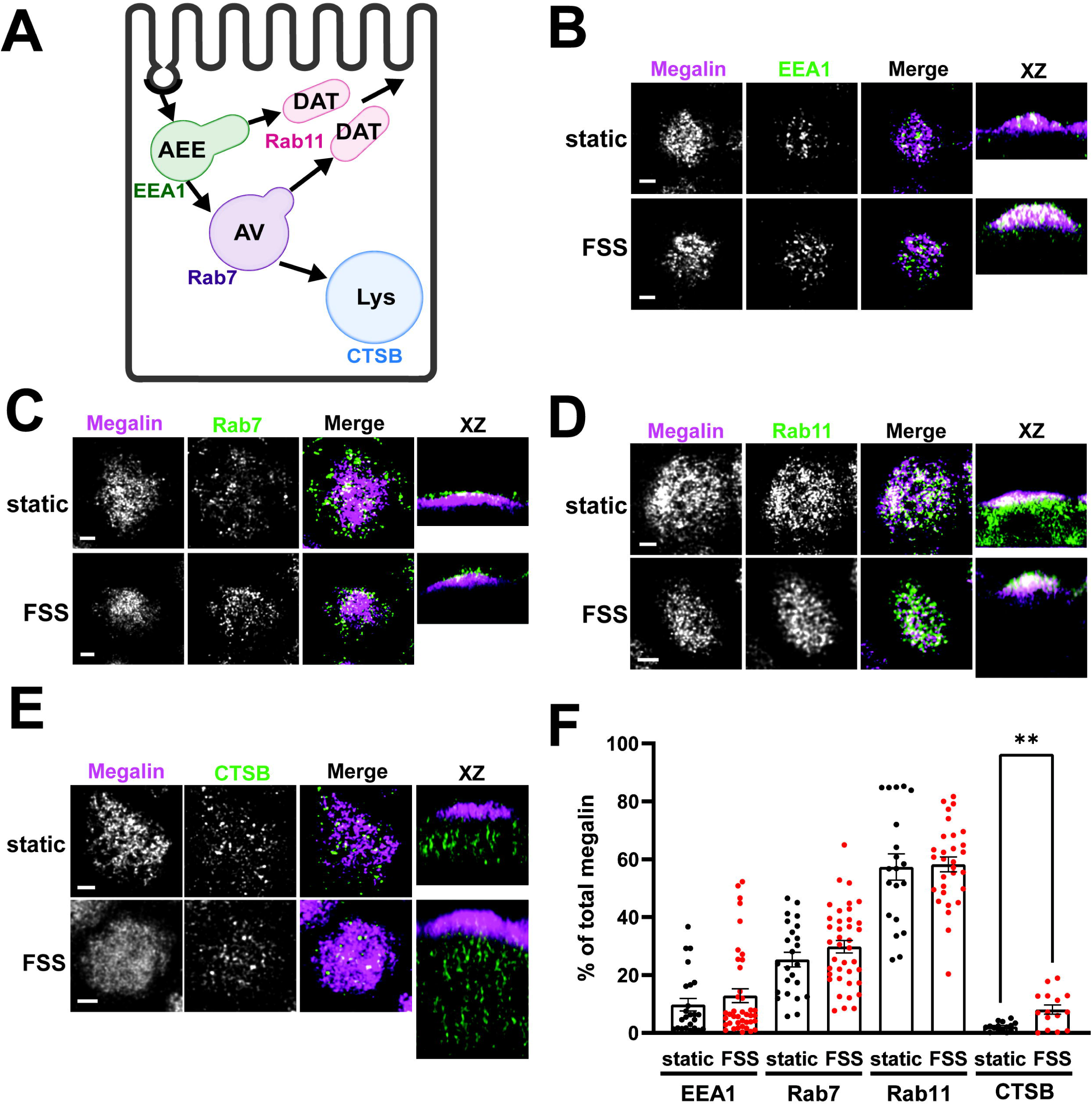
Steady-state distribution of intracellular megalin in static-cultured and FSS-cultured cells is similar. (A) Diagram of the apical endocytic pathway in PT cells with markers used to label each endocytic compartment: apical early endosomes (AEEs) with EEA1, apical vacuoles (AV) with Rab7, dense apical tubules (DATs) with Rab11, lysosomes (Lys) with Cathepsin B (CTSB). Adapted from Shipman et al., 2022, *Function* (Shipman et al., 2022). (B-E) OK cells on permeable supports were fixed and processed to detect megalin colocalization with markers of endocytic compartments in PT cells. Representative sum projection images of 6 planes were cropped to show a region of high colocalization of megalin with (B) EEA1, (C) Rab7, (D) Rab11, and (E) CTSB for both static and FSS cells. XZ representative images are sum projects of the entire image stack. Scale bars: 2 µm. (F) Megalin colocalization with each marker was quantified by Manders’ coefficient over the entire z-stack and plotted as the % of total megalin. Each point represents a single z-stack image. EEA1 P=0.3774; Rab7 P=0.1961; Rab11 P=0.8483; Lys **P=0.0024 by unpaired t-test with Welch’s correction.

To estimate intracellular trafficking rates of megalin in PT cells cultured under static or FSS conditions, we integrated the biochemical and colocalization data into our previously described kinetic model (Shipman et al., 2022). A schematic of the model and associated rate constants is shown in Figure 5A. The colocalization of megalin with each endocytic marker was corrected for the overlap between pairs of markers (Figure S2, Table S3) to estimate the steady state distribution of megalin among endocytic compartment as previously described (Shipman et al., 2022) (Figure 5B). In addition to reduced surface expression, there was a greater fraction of megalin in AEEs and in lysosomes at steady state in FSS-cultured cells. Using the steady state distribution and experimentally measured endocytic rate and surface half-life (Figure 3), we predicted the intracellular trafficking rates in static- and FSS-cultured cells (Figure 5C). The predicted kinetic rates demonstrate faster traffic through all endocytic compartments in FSS-cultured cells. Interestingly, there is slightly less megalin localized in DATs at steady state in FSS-cultured cells, presumably reflecting the ~2-fold more rapid recycling rate in these cells (1% per min). However, this rate is surpassed by the even faster endocytosis of megalin to maintain the low fractional distribution at the apical surface that we measured. Figure 5D tracks the predicted movement through cellular compartments of megalin biotinylated at the apical plasma membrane over a 150 min period. In FSS-cultured cells, megalin is more rapidly removed from the apical surface and reaches a more rapid peak in AEEs. By contrast, in static-cultured cells, endocytosis is slower and megalin accumulates for a longer time in AEEs before trafficking to AVS, DATs and lysosomes.

**Figure 5.**
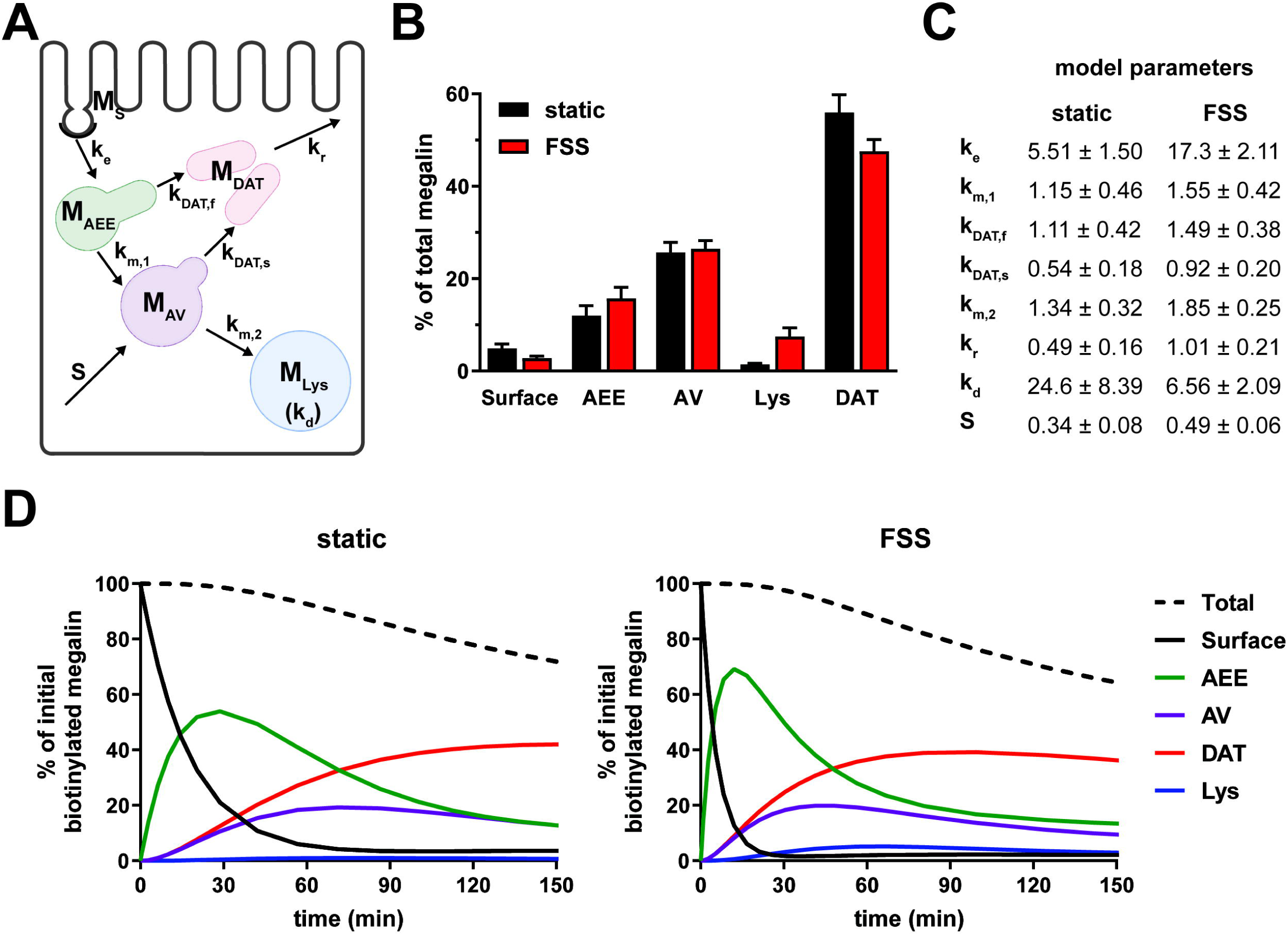
Mathematical model of megalin trafficking reveals an overall increase in trafficking kinetics in FSS-cultured cells with the endocytic rate being the most strongly affected. (A) Graphical representation of the model of megalin traffic modified from Shipman et al., 2022, *Function* (Shipman et al., 2022) depicting the endosomal compartments with coefficients denoting the kinetic rates of megalin traffic between compartments. (B) Megalin steady state distribution among the endocytic compartments, surface, AEEs, AVs, DATs, and lysosomes (Lys) in static-cultured and FSS-cultured cells (mean ± SEM). The distribution was calculated based on the average fractional colocalization of megalin with each marker (Figure 4 and Table S2) and the overlaps between the markers (Figure S2 and Table S3). (C) Kinetic megalin trafficking rates between endosomal compartments were predicted from experimental data for static-cultured and FSS-cultured cells and are presented as % of megalin from the originating compartment trafficked per minute (± SEM). The synthesis (S) rate represents the % of total megalin synthesized per minute and the degradation (k_d_) represents the % of megalin in M_Lys_ that is degraded per minute. (D) The predicted temporal route of megalin biotinylated at the apical surface through each compartment is plotted for static-cultured cells (left) and FSS-cultured cells (right). The synthesis rate is set to zero in this simulation, as no newly biotinylated megalin is created after initial labeling. The total remaining biotinylated megalin over time is plotted as the black dashed lines. Note the more rapid depletion of megalin from the surface and subsequent accumulation in AEEs in FSS cells.

### 3.4 Megalin-driven endocytosis contributes more to gentamicin uptake in cells cultured under shear stress than under static conditions

Understanding the relative contributions of megalin-mediated endocytosis versus other potential internalization pathways to the uptake of a given ligand is important for devising approaches to limit the damage caused by nephrotoxic drugs, including gentamicin and other aminoglycosides. In addition to their uptake via megalin, aminoglycosides can also gain direct access to the cytoplasm via organic transporters (Nagai and Takano, 2014). Uptake of Texas-Red conjugated gentamicin was greater per cell in FSS-cultured cells across a broad range of concentrations (50 nM to 100 nM) compared to static-cultured cells (Figure 6A). To identify the component of uptake that is dependent on megalin, we used receptor associated protein (RAP) as a tool to acutely inhibit megalin-mediated endocytic flux. We previously demonstrated that externally added recombinant RAP acutely and specifically prevents ligand binding to megalin and also silences megalin-driven endocytic flux (Long et al., 2023). In cells with low expression of megalin, treatment with RAP had undetectable effects on endocytic volume (measured using the uptake of fluorescent dextran, which enters cells via clathrin-coated structures used for receptor-mediated endocytosis) (Rbaibi et al., 2023; Long et al., 2023). Addition of RAP to static-cultured cells decreased gentamicin uptake only slightly (Figure 6B). By contrast, RAP significantly decreased gentamicin uptake in FSS-cultured cells by 59% (Figure 6B). The residual uptake of gentamicin in static and FSS-cultured cells was comparable in the presence of RAP, suggesting that other uptake mechanisms operate with similar efficiency in static and FSS-cultured cells.

**Figure 6.**
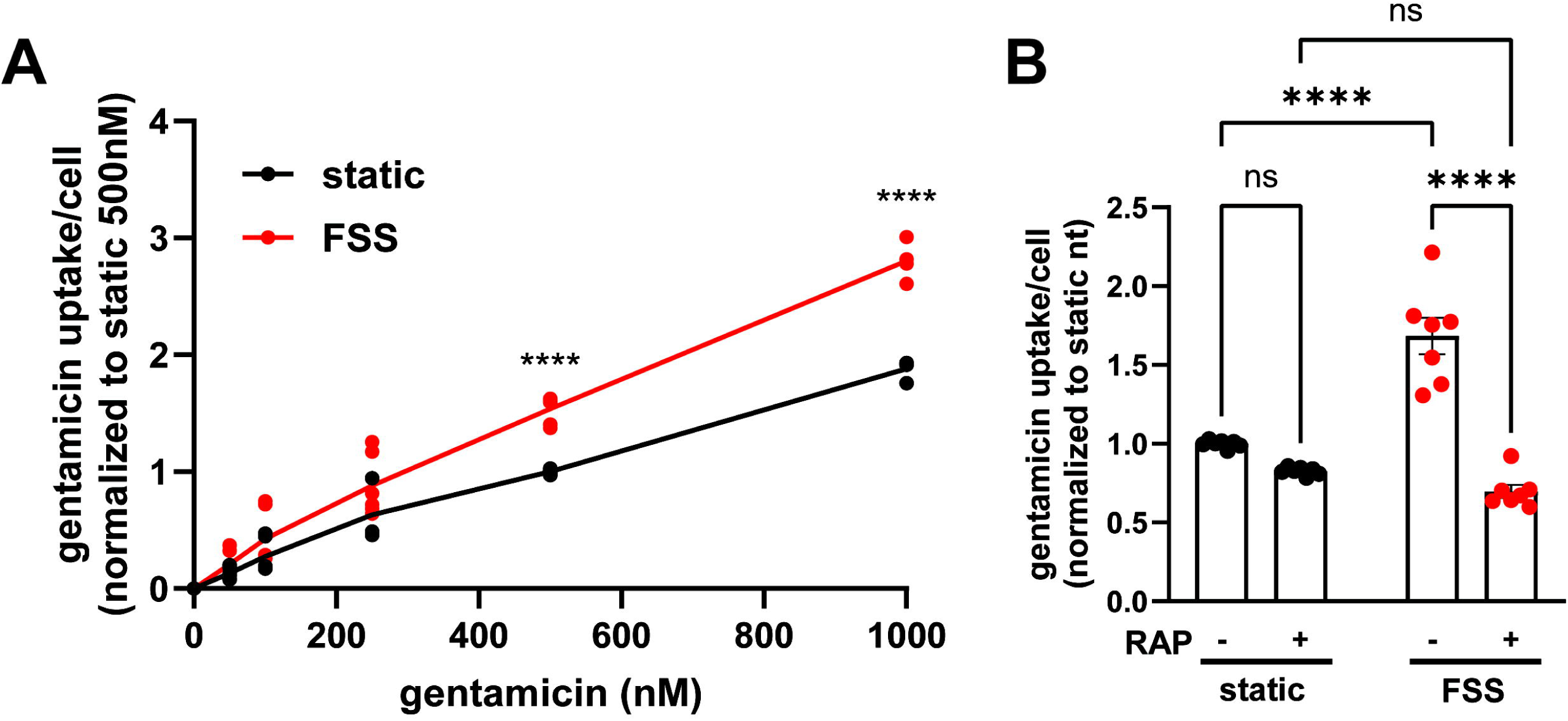
FSS-cultured cells have a larger megalin-driven fraction of apical gentamicin uptake compared to static-cultured cells. (A) OK cells were incubated with 50nM, 100nM, 250nM, 500nM, or 1000nM GTTR added apically for 3 hours. Cells were solubilized and uptake was quantified by spectrofluorimetry per cell. Data is from two to three individual experiments done in duplicate and normalized to static 500nM treated cells. ns=>0.9999, ns=0.9609, ns=0.5514, ns=0.0777, ****P=<0.0001 by Two-way ANOVA with multiple comparisons. (B) OK cells were preincubated with 0.5µM RAP for 30 minutes and then incubated with 500nM GTTR added apically for 3 hours. Cells were solubilized and uptake was quantified by spectrofluorimetry per cell. Data is from three individual experiments done in duplicate or triplicate and normalized to static nontreated. ns=0.1514, ns=0.0583, ****P=<0.0001 by Two-way ANOVA with multiple comparisons.

## 4 Discussion

Our laboratory has previously demonstrated that OK cells cultured under continuous FSS have increased megalin and cubilin expression, greater endocytic uptake, and are morphologically, transcriptionally, and metabolically more representative of the PT *in vivo* compared to static-grown cells (Ren et al., 2019; Long et al., 2017; Park et al., 2020). Here, we showed that changes in the endocytic pathway and more rapid trafficking kinetics in FSS-cultured OK cells also contribute to increased endocytic capacity. These studies begin to define the critical drivers of high apical endocytic capacity in PT cells. Moreover, we demonstrated that internalization of gentamicin via megalin-dependent mechanisms versus other pathways is greatly increased in FSS-cultured cells compared to static conditions. Our data highlight the need to use well-differentiated PT culture models that resemble the PT *in vivo* to develop approaches to limit nephrotoxicity.

By utilizing an approach that integrates biochemical and quantitative imaging assays with mathematical modeling, we determined that megalin trafficking at each step of the endocytic pathway is faster in FSS-cultured OK cells compared to cells cultured under static conditions. The endocytic rate of megalin was the most affected and was three times greater in FSS-cultured cells compared with static-grown cells. Consequently, the steady state fraction of megalin present at the apical surface is lower in FSS-cultured cells compared with control cells. At normally filtered concentrations, albumin (40 µg/mL) binds primarily (approximately 90%) to cubilin (Ren et al., 2020), while megalin expression drives overall endocytic flux (Long et al., 2023). Thus, this substantial increase in endocytic rate, together with the 3.8-fold greater expression of megalin and 2.3-fold greater expression of cubilin, drives the >5-fold increase in endocytic capacity for albumin uptake in FSS-grown cells.

Coincident with the considerable increase in the endocytic rate, transmission electron microscopy revealed an expansion in endocytic compartments filling the subapical cytoplasm (Long et al., 2017). Here, we examined the distribution of intracellular megalin across the endocytic compartments. Surprisingly, the fractional colocalization of megalin with markers of AEEs (EEA1), AVs (Rab7), and DATs (Rab11a) was not different between static- and FSS-cultured cells. However, there was a notable change in the distribution of Rab11a which was distinctly more concentrated within the subapical region of FSS-cultured cells. In addition, we have previously reported a significant increase in the expression of Rab11a in FSS-cultured cells (Long et al., 2017). This increase in expression and polarized distribution of Rab11a may contribute to the more rapid (2-fold) recycling rate of megalin predicted by our kinetic model.

Our kinetic model predicts that the fraction of megalin that is recycled versus targeted for degradation is greater in FSS-cultured cells compared with static-cultured cells. At first glance, this is perhaps surprising, given that surface-tagged megalin has a shorter half-life and greater colocalization with LAMP1 in FSS-cultured versus static-cultured cells. However, this is a consequence of the overall more rapid traffic in FSS-cultured compared to static-cultured cells. Despite its shorter half-life, megalin undergoes more rounds of endocytosis and recycling during its lifespan.

Another surprising finding in our study is the differential changes in the expression and distribution of cubilin versus megalin in FSS-cultured compared to static-cultured cells. Previous biochemical and morphological studies have suggested that, in the PT, cubilin and megalin traffic as a complex (Moestrup et al., 1998; Ahuja et al., 2008). We also showed that megalin- and cubilin-mediated uptake of albumin is coordinately altered in our cell culture model of Dent disease (Shipman et al., 2023). By contrast, here we found a larger increase in the total expression of megalin compared to cubilin (3.8-fold vs. 2.3-fold, respectively) in FSS-cultured versus static-cultured cells, along with a greater decrease in the fractional surface localization of cubilin compared to megalin (60% vs. 43%). Megalin was recently shown to assemble as a homodimer, whereas CUBAM contains three cubilin molecules (Beenken et al., 2023; Larsen et al., 2018). This could contribute to discordant changes in megalin and cubilin distribution as a similar fractional change in receptor complexes at the surface would result in a 1.5x larger change in the number of cubilin molecules compared to megalin. However, to be able to precisely correlate changes in receptor expression and traffic with ligand uptake would require determination of both the endocytic rate of cubilin and the stoichiometry of albumin binding to these complexes. Regardless, our studies are consistent with our prior observations that megalin and cubilin continue to internalize ligands when expressed independently of each other (Long et al., 2022; Rbaibi et al., 2023; Long et al., 2023).

Many current *in vitro* models of drug-induced nephrotoxicity are poorly predictive of toxicity in humans (Soo et al., 2018). Our findings highlight the importance of using a well differentiated cell model to study drug uptake by the PT. We found that, contrary to static-cultured OK cells, apical uptake of gentamicin in FSS-cultured cells is primarily megalin driven, and that the increased megalin expression and faster endocytic rate contributes to the increased gentamicin accumulation. Consistent with this, Molitoris and colleagues recently demonstrated that infusion of RAP significantly reduced the PT accumulation and nephrotoxic effects of gentamicin in rats (Wagner et al., 2023). Gentamicin can also enter PT cells basolaterally via cation transporters such as TRPV4 (Nagai and Takano, 2014), although the contribution of gentamicin uptake via this pathway was not specifically examined. Accumulation of reactive oxygen species that impair metabolism and trigger pathologic signaling is thought to underlie the toxic effects of aminoglycosides. The similarities in endocytic capacity and metabolism between FSS-cultured cells and the PT *in vivo* highlight this model as a tool to assess the uptake pathways of clinically important drugs as well as the effects of potential therapeutics that limit nephrotoxicity.

## 6 Author Contributions

O.A.W. and K.E.S conceived and designed research; E.M.L., K.R.L., I.A.C, and K.E.S. performed experiments; E.M.L. and K.E.S. analyzed data; O.A.W., E.M.L. and K.E.S. interpreted results of experiments; E.M.L. and K.E.S. prepared figures; E.M.L. and K.E.S. drafted manuscript; E.M.L., K.E.S., and O.A.W. edited and revised manuscript; K.R.L., E.M.L., K.E.S., I.A.C., and O.A.W. approved final version of manuscript.

## Acknowledgments

We thank Drs. Daniel Biemesderfer and Peter Aronson (Yale University) for providing the anti-megalin antibody.

