## Supplemental Material for "Fluid Shear Stress-Induced Changes in Megalin Trafficking Enhance Endocytic Capacity in Proximal Tubule Cells"

### Supplemental Materials

#### Table S1. Antibodies for indirect immunofluorescence in OK cells

| Primary Antibody Target or Secondary | Company | Catalog # | Source | Working Dilution |
|---|---|---|---|---|
| Megalin | (Zou et al. 2004) | -- | Rabbit | 1:1000 |
| Rab11a | Abcam | ab65200 | Rabbit | 1:500 |
| Rab7 | Sigma | R8779 | Mouse | 1:200 |
| Cathepsin B | Cell Signaling | 31718 | Rabbit | 1:500 |
| EEA1 (E-8) | Santa Cruz | sc-365652 | Mouse | 1:50 |
| Anti-mouse Alexa Fluor 488 | Invitrogen | A11029 | Goat | 1:500 |
| Anti-rabbit Alexa Fluor 488 | Invitrogen | A11034 | Goat | 1:500 |
| Anti-mouse Alexa Fluor 647 | Invitrogen | A21236 | Goat | 1:500 |
| Anti-mouse Alexa Fluor 647 | Invitrogen | A21245 | Goat | 1:500 |
| F(ab) anti-mouse Alexa Fluor 488 | Jackson Immuno | 115-547-003 | Goat | 1:500 |
| F(ab) anti-rabbit Alexa Fluor 488 | Jackson Immuno | 111-547-003 | Goat | 1:500 |
| F(ab) anti-mouse (unconjugated) | Jackson Immuno | 115-007-003 | Goat | 1:65 |
| F(ab) anti-rabbit (unconjugated) | Jackson Immuno | 111-007-003 | Goat | 1:65 |

#### Table S2. Average fractional colocalization (Manders' coefficient) of megalin with endocytic markers

| | static | | | FSS | | |
|---|---|---|---|---|---|---|
| **Manders' Coefficient** | **Mean** | **SEM** | **N** | **Mean** | **SEM** | **N** |
| $M_{EEA1}$ | 0.0975 | 0.0216 | 24 | 0.1287 | 0.0238 | 40 |
| $M_{Rab7}$ | 0.2533 | 0.0253 | 24 | 0.2979 | 0.0218 | 39 |
| $M_{Rab11}$ | 0.5731 | 0.0454 | 20 | 0.5824 | 0.0255 | 30 |
| $M_{CTSB}$ | 0.0213 | 0.0040 | 15 | 0.0807 | 0.0160 | 15 |

**Table S3. Average fractional overlap (Manders' coefficient) between endocytic markers**

| | static | | | FSS | | |
|---|---|---|---|---|---|---|
| **Manders' Coefficient** | **Mean** | **SEM** | **N** | **Mean** | **SEM** | **N** |
| $EEA1_{Rab7}$ | 0.5073 | 0.0555 | 29 | 0.5583 | 0.0436 | 49 |
| $EEA1_{Rab11}$ | 0.2539 | 0.0194 | 20 | 0.2970 | 0.0197 | 35 |
| $Rab7_{EEA1}$ | 0.1587 | 0.0235 | 29 | 0.2513 | 0.0253 | 49 |
| $Rab7_{Rab11}$ | 0.2602 | 0.0429 | 25 | 0.4466 | 0.0333 | 35 |
| $Rab7_{CTSB}$ | 0.0180 | 0.0026 | 10 | 0.0217 | 0.0048 | 10 |
| $Rab11_{EEA1}$ | 0.0663 | 0.0077 | 20 | 0.0792 | 0.0191 | 35 |
| $Rab11_{Rab7}$ | 0.1415 | 0.0248 | 25 | 0.2528 | 0.0260 | 35 |
| $Rab11_{CTSB}$ | 0.0164 | 0.0054 | 10 | 0.0193 | 0.0088 | 9 |
| $CTSB_{Rab7}$ | 0.3487 | 0.0297 | 10 | 0.0633 | 0.0093 | 10 |
| $CTSB_{Rab11}$ | 0.4652 | 0.0534 | 10 | 0.2443 | 0.0563 | 9 |

### Supplemental Figure Legends

**Figure S1. Quantification of cell heights in static- and FSS-cultured cells.** (A) OK cells on permeable supports were fixed and stained for megalin, Rab11a and actin. To determine cell height, using ImageJ software and XZ stacks, line objects were drawn from the base of the cell to the tip of the microvilli based on actin staining. Each dot represents the height of one cell. ****P=<0.0001 by unpaired t-test, n=85-86 cells. (B-C) Representative images of actin, megalin, and Rab11 staining in static-cultured (B) and FSS-cultured (C) cells. XZ images are sum projections of the entire cell stack. Scale bars: 2μm.

**Figure S2. Quantification of the overlap between endocytic markers in static- and FSS-cultured cells.** (A-C) OK cells on permeable supports were fixed and processed to detect colocalization between the indicated pairs of endocytic compartment markers. Quantification by Manders' coefficient over the entire z-stack, shown as the % of total for the following: (A) EEA1 and Rab7, (B) Rab7 and Rab11, and (C) EEA1 and Rab11. Each point represents a single z-stack image. *P=0.0345, ***P=0.0002 by Two-way ANOVA with multiple comparisons.

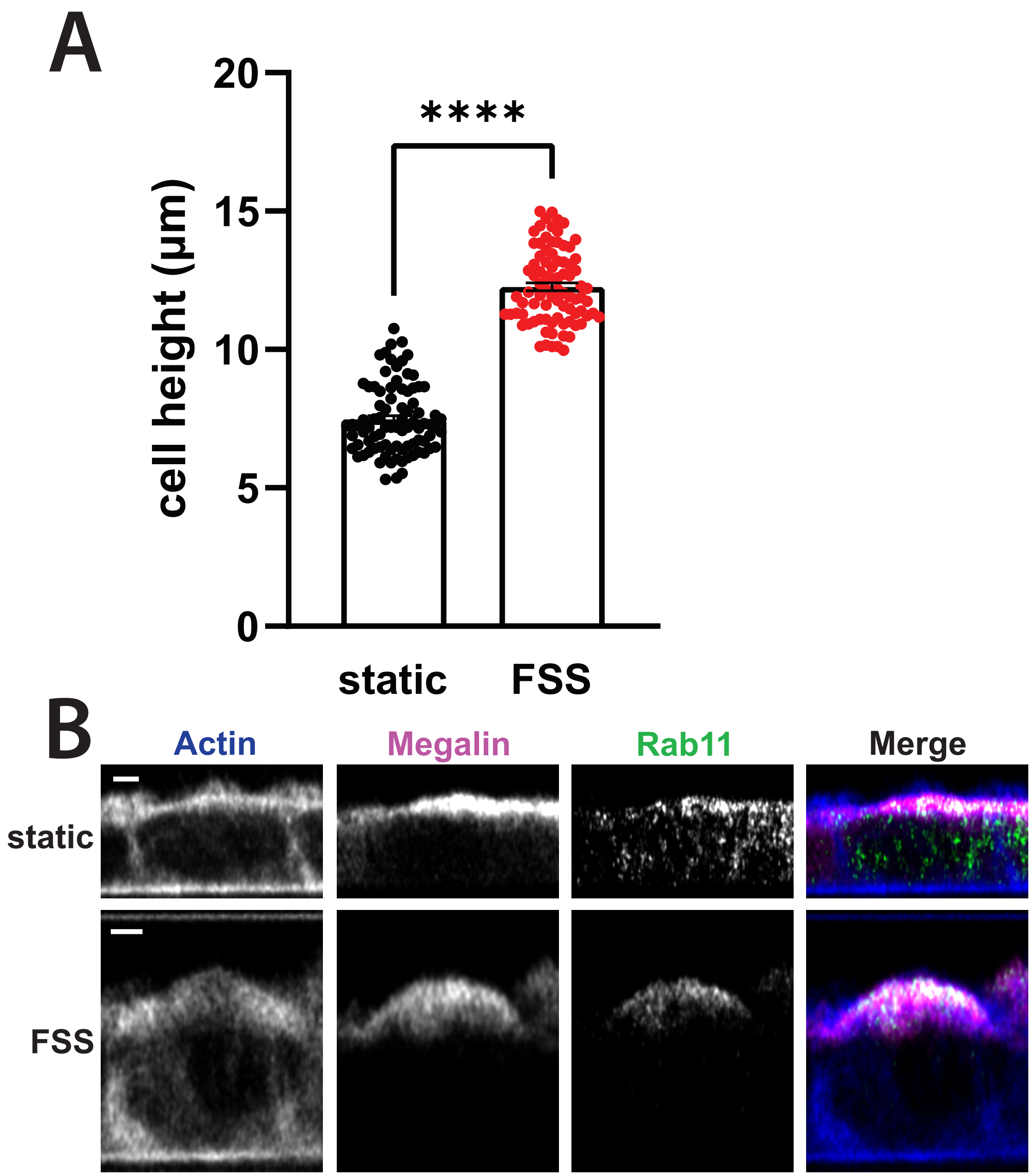
A
20
15
10
5
0
cell height (µm)
****
static
FSS
B
Actin
Megalin
Rab11
Merge
static
FSS

Lackner et al. Figure S2

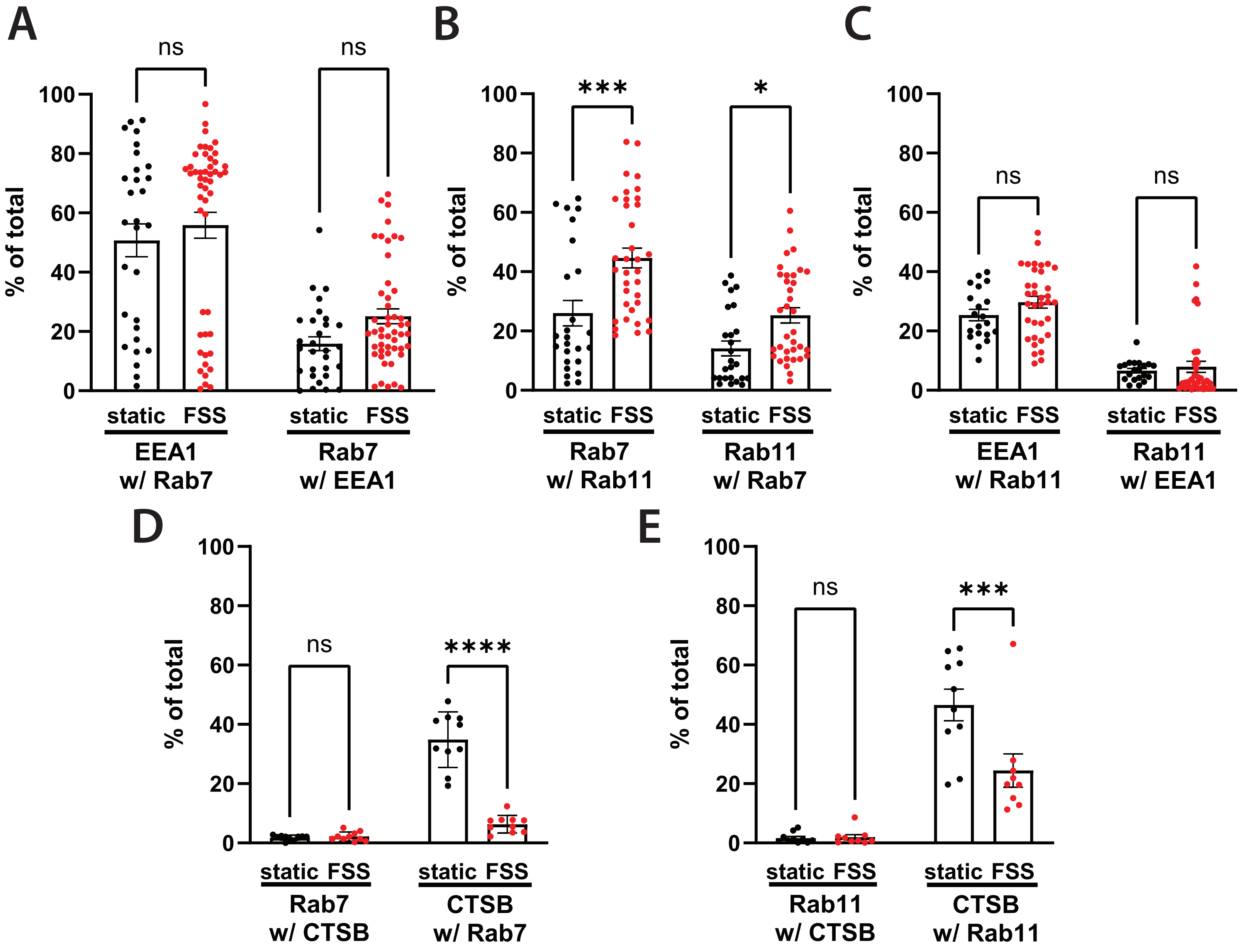
